# CRK2 modulates flowering in *Arabidopsis* together with GLYCINE- RICH RNA BINDING PROTEIN 7 (GRP7)

**DOI:** 10.1101/2025.08.05.668606

**Authors:** Francisco J. Colina, Julia Krasensky-Wrzaczek, Alegría Pérez-Guillén, Michael Wrzaczek

**Affiliations:** Institute of Plant Molecular Biology, Biology Centre, Czech Academy of Sciences, 370 05 České Budějovice, Czech Republic; Faculty of Science, University of South Bohemia, 370 05 České Budějovice, Czech Republic; Estación Experimental del Zaidín, Consejo Superior de Investigaciones Científicas (EEZ-CSIC), 18008 Granada, Spain

**Keywords:** Arabidopsis, CRK2, GRP7, SRBP1, gibberellin (GA), flowering

## Abstract

Timely flowering is critical for the reproductive success of plants. Plants adjust flowering time through integration of multiple internal and environmental cues. A complex signalling system lies behind this integration, including multiple proteins like the GLYCINE-RICH RNA-BINDING PROTEIN 7 (GRP7), linked to the modulation of stress and flowering in Arabidopsis. Available data show the interaction between GRP7 and the receptor-like kinase CYSTEINE-RICH RECEPTOR-LIKE KINASE 2 (CRK2). Like GRP7, CRK2 plays multifaceted roles in the coordination of stress responses and development, but its role in development remains unclear. Phenotypic analyses of the *crk2* and *grp7-1* single mutants as well as the *crk2/grp7-1* double mutant supported the interaction between CRK2 and GRP7. Moreover, exogenous gibberellic acid did not recover the rosette phenotype of *crk2* mutants but as in sensitive plants, inhibited GA-biosynthesis transcripts. The pseudo insensitivity of *crk2* mutant was recovered in the *crk2/grp7-1* line pointing to a common role of CRK2 and GRP7 in the modulation of gibberellin signalling. Together, CRK2 and GRP7 integrate hormonal and spatial signals required for floral transition. These insights highlight a previously unrecognized layer of regulation in GA signalling and open new avenues to understand how membrane-localized kinases and RNA-binding proteins intersect to control developmental timing.

## Introduction

In this article, we explored how the interaction with the GLYCINE-RICH RNA-BINDING PROTEIN 7 (GRP7) may allow CYSTEINE-RICH RECEPTOR-LIKE KINASE 2 (CRK2) to modulate flowering time in Arabidopsis. Our model points to a functional interaction between CRK2 and GRP7 that allows the RLK to introduce inputs into signalling mechanisms which, mediated by the plant phytohormone gibberellic acid (GA), might modulate flowering and environmental responses in Arabidopsis.

Timely flowering is crucial for plants, as it ensures production of viable seeds and thereby the next generation. Plants adjust flowering time through the integration of internal and environmental cues, including energy status, temperature and photoperiod to optimize reproductive output in a changing environment (Andrés & Coupland, 2012; Wahl *et al*., 2013; Song *et al*., 2015; Yu *et al*., 2015; Kazan & Lyons, 2016). This regulation involves pathways responsive to light (photoperiod) and temperature (vernalization) as well as endogenous signals (autonomous pathway, sugar, age). Among these, GA plays central roles in flowering control (Langridge, 1957; Wilson *et al*., 1992; Porri *et al*., 2012; Mateos *et al*., 2015; Bao *et al*., 2020), but also have roles in environmental responses (Colebrook *et al*., 2014). Despite extensive knowledge of the core flowering pathways, the full molecular network that coordinates them and integrates flowering with other responses to the environment remains incomplete (Kazan & Lyons, 2016).

Receptor-like kinases (RLKs) are emerging as integrators of developmental transitions, including flowering, and environmental stimuli (Wang *et al*., 2020). Among them, CRK2 plays multifaceted roles in stress signalling (Hunter *et al*., 2019; Kimura *et al*., 2020) and development, as suggested by the late flowering and small rosette size of the *crk2* mutant (Bourdais *et al*., 2015). Interestingly, CRK2 has previously been associated with GA signalling (Cao *et al*., 2006; Bassel *et al*., 2011). However, the molecular mechanism of the integration of CRK2 with GA signalling has so far not been elucidated.

GRP7, also known as SMALL RNA-BINDING PROTEIN 1 (SRBP1), a diurnally modulated RNA-binding protein (Staiger *et al*., 2003), has been identified as an *in planta* interaction partner of CRK2 (Hunter *et al*., 2019). Like CRK2, GRP7 modulates stress responses and developmental processes, including flowering time (Nicaise *et al*., 2013; Wang *et al*., 2020). GRP7 negatively regulates its own expression (Staiger *et al*., 2003) as well as the expression of the central flowering repressor *FLOWERING LOCUS C* (*FLC*) (Streitner *et al*., 2008; Steffen *et al*., 2019) within the autonomous pathway (AP). Moreover, phenotypes of GRP7 overexpression plants suggest a role as negative modulator of GA signalling (Löhr *et al*., 2014). By participating in AP and GA signalling, GRP7 provides a possible mechanism for its partner, CRK2 to input signals into the GA and/or AP flowering pathways.

## Results and discussion

### CRK2 modulates development in Arabidopsis through endogenous pathways

Under long day (LD) conditions, *crk2* exhibits a late flowering phenotype (Bourdais *et al*., 2015). Interestingly, *crk2* showed an even more severe flowering delay in short day (SD) (Figure 1a; Figure S1; Table S1; Table S2a). In SD-grown Arabidopsis, flowering transition relies on endogenous pathways, including the AP and GA pathways. Mutants in these pathways, frequently show a more severe flowering delay in SD compared to LD (Macknight *et al*., 2002; Ausín *et al*., 2004; Streitner *et al*., 2008; Steffen *et al*., 2019), suggesting that CRK2 might act via similar mechanisms. Notably, *crk2* plants also had fewer rosette leaves compared to Col-0 at the time of bolting (Figure 1b, Table S2b). Similar phenotypes where flowering time does not correlate with leaf number are rare (Koornneef *et al*., 1991; Méndez-Vigo *et al*., 2010).

**Figure 1.**
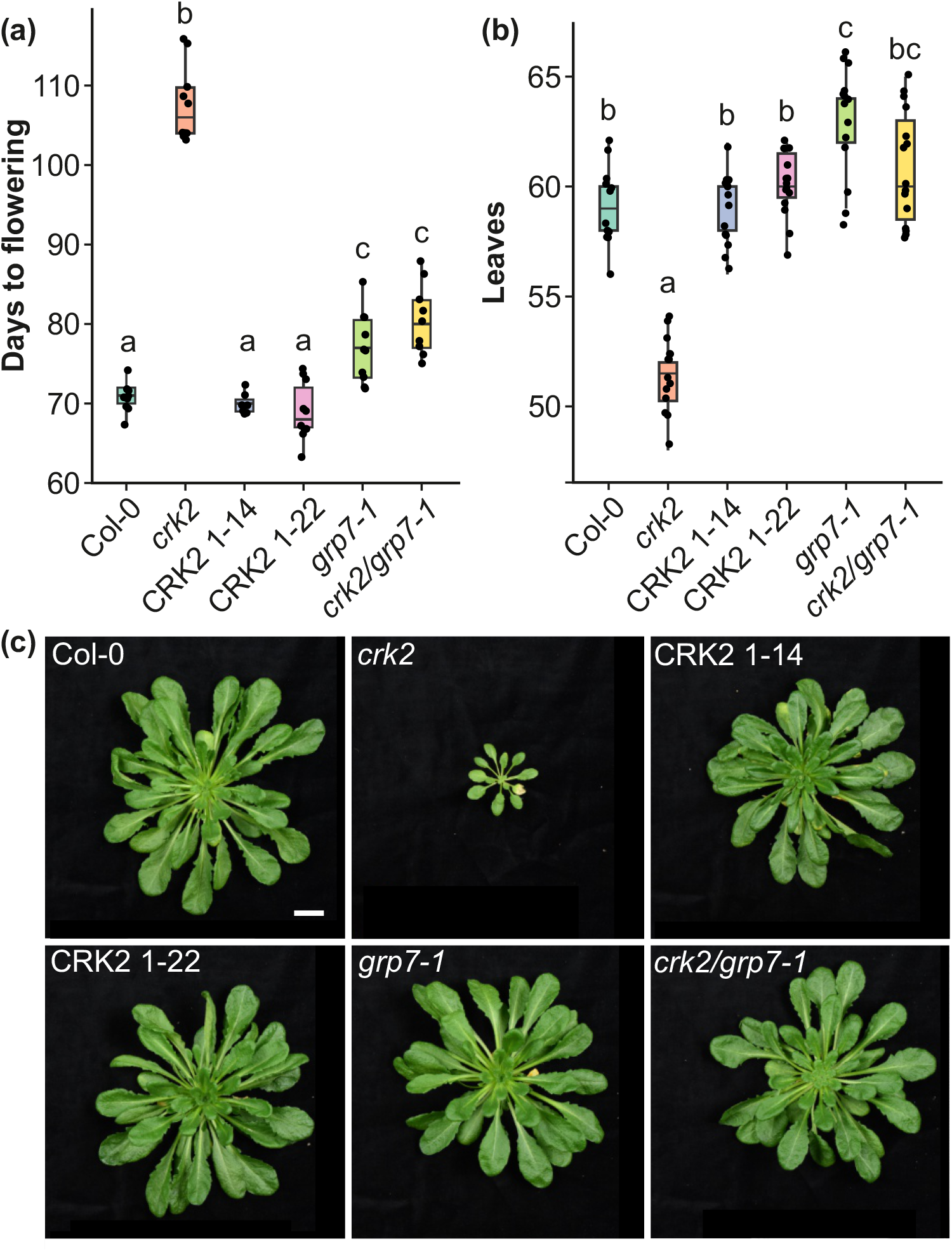
CRK2 and GRP7 modulate Arabidopsis flowering time under non inductive conditions. **(a)** Flowering time (in days) of *Arabidopsis thaliana* Col-0, *crk2, grp7-1, crk2/grp7-1* double mutant, and two complementation lines (*CRK2p::CRK2YFP/crk2* #1-14 and #1-22) grown under short-day (SD) conditions. Statistical significance was assessed by Kruskal–Wallis followed by Wilcoxon post hoc test. Letters indicate statistically significant differences between genotypes (p < 0.05). **(b)** Number of rosette leaves at flowering for the same genotypes under SD conditions. Statistical significance was assessed by one-way ANOVA followed by Tukey’s post hoc test. Letters indicate statistically significant differences between genotypes (p < 0.05). **(c)** Representative rosettes of 75-day-old plants grown under SD conditions. **(a, b)** Data are presented as boxplots showing the median, interquartile range (IQR), and whiskers extending to 1.5×IQR. Individual data points are overlaid.

### CRK2 interacts functionally with the RNA binding protein GRP7

Interestingly, under SD *crk2* flowered even later than *grp7-1* and *crk2/grp7-1*, which flowered later than Col-0 (Figure 1a, Table S2a). As previously described, the rosette size of *crk2* was small compared to Col-0 (Kimura *et al*., 2020) while rosettes of *grp7-1* and *crk2/grp7-1* showed a similar size to Col-0 (Figure 1c). This partial rescue of the *crk2* developmental phenotypes by *grp7-1* adds to previous evidence on CRK2-GRP7 interaction (Hunter *et al*., 2019), suggesting that GRP7 and CRK2 interact functionally. This links CRK2 with the GA and AP pathways, which are both modulated by GRP7 (Kim *et al*., 2008; Löhr *et al*., 2014; Xiao *et al*., 2015; Wang *et al*., 2020).

### CRK2 modulates GA signalling with the involvement of GRP7

Both CRK2 and GRP7 are linked to GA biosynthesis and signalling (Cao *et al*., 2006; Kim *et al*., 2008; Bassel *et al*., 2011; Löhr *et al*., 2014; Wang *et al*., 2020). Exogenous GA treatment reduces *CRK2* transcript abundance in the GA biosynthesis deficient *ga1-3* mutant (Bassel *et al*., 2011). Like *crk2*, GA insensitive mutants like *gai* display small rosette size and a severe flowering delay under SD (Wilson *et al*., 1992). The *gai* mutant carries a GA-resistant allele of the GAI DELLA, a repressor of GA responses degraded upon GA perception (Peng *et al*., 1997). Interestingly, DELLAs positively regulate *CRK2* transcript levels (Cao *et al*., 2006). However, whether CRK2 regulates DELLAs is an open question, which can be tested in the future by evaluating DELLA stability in *crk2*. Besides the CRK2 link with GA signalling, *CRK2* transcript abundance also correlates with the GA biosynthesis-related transcript *GIBBERELLIN 3-BETA-DIOXYGENASE 1* (*GA3ox1*) (Bassel *et al*., 2011). GRP7 overexpression plants are sensitive to exogenous GA but contain reduced levels of bioactive GAs and reduced transcript abundance of GA biosynthesis and response genes like *ENT-KAURENE SYNTHASE* (*GA2*; GA biosynthesis) and *GIBBERELLIC ACID STIMULATED ARABIDOPSIS 9* (*GASA9*; GA response) (Löhr *et al*., 2014).

To evaluate whether CRK2 and GRP7 genetically interact in the GA pathway we investigated the impact of GA treatment on *crk2, grp7-1*, the *crk2/grp7-1* double mutant, and *crk2* complementation lines (Kimura *et al*., 2020) in comparison to Col-0. As previously described, treatment with GA increased the rosette size of Col-0 compared to mock treatment (Figure 2a) (Ribeiro *et al*., 2012; Löhr *et al*., 2014) while *crk2* was insensitive to the phytohormone (Figure 2a). By contrast, GA treatment increased rosette size in *grp7-1* and *crk2/grp7-1*, similar to Col-0 (Figure 2a), suggesting a functional rescue of the GA sensitivity by *grp7-1*. In addition, *crk2* showed higher abundance of GA biosynthesis related *GA2* and *GA3ox1* transcripts compared to Col-0 under mock conditions (Figure 2b Figure S2, Table S3a, Table S4). GA3ox1, involved in the final steps of bioactive GA synthesis, is subject to feedback repression by GA (Chiang *et al*., 1995). Surprisingly, while GA treatment did not affect *crk2* rosette size, transcript abundance of *GA3ox1* was reduced following GA treatment in *crk2* similar to Col-0 (Figure 2b; Table S3a). The change in *GA3ox1* abundance in *crk2* showed functionally negative feedback and suggested that in *crk2*, GA signalling is impaired downstream of perception. Transcript abundance of *GA3ox1* in *grp7-1* and *crk2/grp7-1* was similar to Col-0 under mock conditions. In our experiment, all included genotypes but *crk2* and *grp7-1* showed a downward trend in the abundance of the GA-response transcript *GASA9* (Figure 2c; Table S3b). Addition of GA reduces *GASA9* abundance in Col-0 (Zhang & Wang, 2008), suggesting altered transcriptional responses to GA in *crk2* and *grp7-1*.

**Figure 2.**
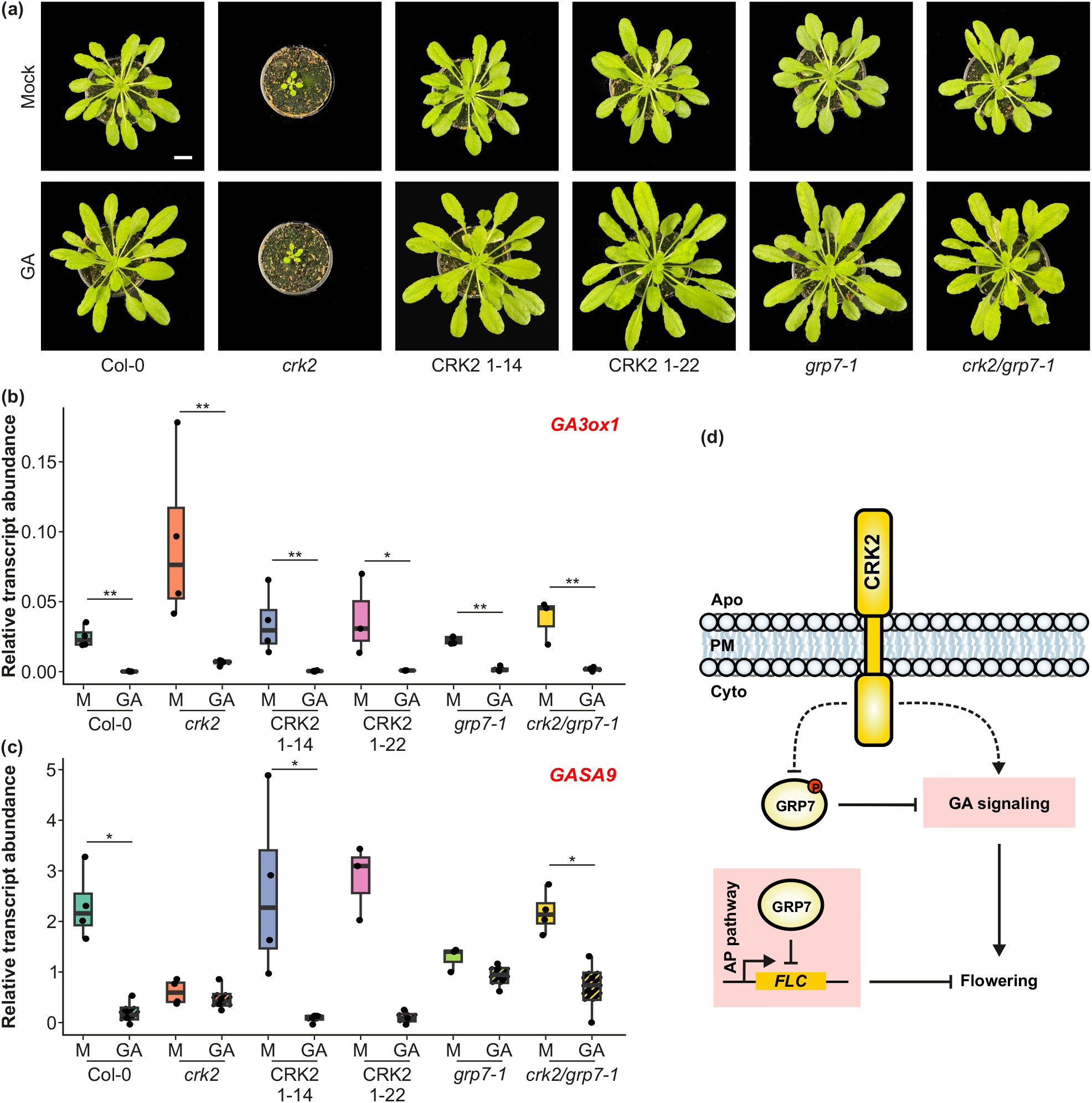
Effect of GA treatment. **(a)** Representative rosettes of two-month-old plants grown under SD conditions and under weekly spraying with mock or GA_3_ containing solutions. **(b)** Abundance of GA-biosynthesis transcript *GA3ox1*. Pairwise comparisons were performed within each genotype using Wilcoxon rank-sum test. P-values were adjusted for multiple testing using the Benjamini– Hochberg (BH) method (* p<0.1, ** p<0.05). **(c)** Abundance of GA-responsive transcript *GASA9*. Pairwise comparisons were performed within each genotype using Wilcoxon rank-sum test. P-values were adjusted for multiple testing using the Benjamini–Hochberg (BH) method (* p<0.1). **(b, c)** Data are presented as boxplots showing the median, interquartile range (IQR), and whiskers extending to 1.5×IQR. Individual data points are overlaid. **(d)** Hypothetical model for the CRK2-GRP7-based modulation of flowering time in Arabidopsis. CRK2 is likely a flowering modulator in Arabidopsis. Our data points to a functional interaction between CRK2 and GRP7 focused on GA signalling and independent of the roles of GRP7 in the modulation of *FLC*. In our hypothesis, the RLK acts over the GA signalling pathway via phosphorylation-based modulation (likely inhibition) of GRP7 but might also exhibit a GRP7-independent influence on GA signalling.

The phenotype of the *crk2*/*grp7-1* double mutant indicates a genetic interaction between CRK2 and GRP7 in GA signalling. However, whether they act in parallel or at least partially through the same pathway is still an open question. GRP7 likely is a phosphorylation substrate for CRK2 (Figure 2d), which carries a functional kinase domain (Hunter *et al*., 2019; Kimura *et al*., 2020). The recombinant cytosolic region of CRK2 (CRK2cyt) phosphorylates recombinant GRP7, but not the GRP7 interactor JACALIN-LECTIN LIKE1 (JAC1), *in vitro* (Figure S3). Moreover, previous works found that the RLK FERONIA (FER) phosphorylates GRP7 in its RGG domain (Wang *et al*., 2020). This phosphorylation impacts GRP7 subcellular location and liquid-liquid phase separation (LLPS) ability which in turn impacts its affinity for RNA (Wang *et al*., 2020; Xu *et al*., 2024; Lühmann *et al*., 2024). Phosphorylation-mediated changes in GRP7 likely impact the splicing-mediated maturation of stress- and development-related transcripts (Wang *et al*., 2020). If GRP7 is phosphorylated by CRK2 *in vivo* it would be thus intriguing to describe the phosphorylation sites and their relevance in the presented genetic dependency between CRK2 and GRP7 at GA signalling. Interestingly, available RNA immunoprecipitation data shows that GRP7 targets multiple GA biosynthesis and response (*GASA*) transcripts (Meyer *et al*., 2017; Wang *et al*., 2020) suggesting a direct effect of GRP7 on the maturation of GA response transcripts and providing a promising starting point for future approaches. It will be thus interesting to analyze whether CRK2, potentially through phosphorylation of GRP7, modulates the interaction of GRP7 with GA-related transcripts and their processing. Using *in vivo* phosphoproteomic analyses of Col-0 and *crk2* and RNA immunoprecipitation (RIP)-seq of plants expressing non-phosphorylatable variants of GRP7 could provide these insights in the future.

Despite a possible direct role of CRK2 on the modulation of GA signalling via GRP7, the functional interaction between CRK2 and GRP7 here described also provides potential links between CRK2 and the AP. The AP converges with the cold-associated vernalization flowering pathway at the repression of *FLC*. Therefore, the flowering delay of AP mutants like *grp7-1* is reversed by cold treatment (Streitner *et al*., 2008; Steffen *et al*., 2019). Interestingly, vernalization at 4ºC for 45 days rescued the flowering delay of *grp7-1* and *crk2/grp7-1*, but only partially for *crk2* (Figure S4a; Table S5. Despite the vernalization effect, Col-0 and *crk2* showed a similar abundance of *FLC* (Figure S4b, c; Table S6) and vernalization reduced *FLC* transcript levels similarly in Col-0 and *crk2* (Figure S4c; Table S6c). By contrast, the *crk2/grp7-1* double mutant showed high *FLC* abundance similar to *grp7-1* (Figure S4a; Table S6a). Together with the similar flowering phenotypes of *grp7-1* and *crk2/grp7-1* (Figure 1; Table S2, S6) this suggests that GRP7 modulates *FLC* independently of CRK2 (Figure 2d). Despite this intriguing finding, future works should determine whether the identified lack of effect on *FLC* in *crk2* at the whole-plant level is explained by a targeted effect of GRP7 phosphorylation on a specific subset of its targets or is associated with a spatial functional compartmentalization of the CRK2-GRP7 interaction.

### Are CRK2 and GRP7 involved in the modulation of symplastic transport of flowering signals?

Despite the many open questions on how CRK2 contributes to flowering via GA and whether GRP7 is involved, their known roles may contribute to lead future approaches towards characterizing these mechanisms. CRK2 has previously been linked to regulation of callose deposition (Hunter *et al*., 2019). Callose has important roles in controlling plasmodesmal connectivity and contributes to regulating transport of mobile protein flowering signals like the flowering promotor FLOWERING LOCUS T (FT) to the SAM (Murata *et al*., 2025). Interestingly, altered callose deposition has been observed in *grp7-1* after pathogen infection (Fu *et al*., 2007). Therefore, CRK2 and GRP7 may modulate symplastic transport via callose deposition. Moreover, GRP7 has been linked to the transport of short RNAs (sRNAs) via plasmodesmata (Yan *et al*., 2020), and has been identified in phloem exudates (Batailler *et al*., 2012) and plasmodesmata enriched fractions (Fernandez-Calvino *et al*., 2011), suggesting that GRP7 might be a cell-to-cell mobile protein. Mobile GRP7 could be able to shuttle its RNA load between cells. RNA transport between cells, especially when this transport involves long-distance movement via phloem, is currently matter of active debate following recent reanalysis of mobile RNA datasets, which showed that RNA transport is likely less common than previously suggested (Paajanen *et al*., 2025). However, targeted approaches have successfully identified the mRNA of the DELLA protein *GAI*, to move through the phloem and induce dwarfism in distal tissues (Haywood *et al*., 2005). It is thus tempting to speculate on a role of CRK2 and GRP7 in the modulation of the transport and/or the maturation of GA-related transcripts. CRK2 might modulate GRP7 function to coordinate RNA processing and intercellular transport, thereby integrating hormonal, spatial, and post-transcriptional regulatory layers essential for developmental transitions such as flowering. In summary, our results position CRK2 as a key modulator of GA signalling, acting partly through GRP7 to control development (Fig. 2d).

## Acknowledgments

The authors would like to thank Sunita Jindal, María Carbó and Adam Zeiner (**Institute of Plant Molecular Biology, Biology Centre, Czech Academy of Sciences, Czechia**), and to Ainhoa Martínez Medina, Iván Fernández López and Ana María Jiménez Jiménez (**Estación Experimental del Zaidín, Consejo Superior de Investigaciones Científicas, Spain**) for critical comments on the article. This work was supported by the Czech Academy of Sciences (startup grant to MW) and by the Czech Science Foundation (grant 23-04866S). FC was funded by EMBO (award number: ALTF 1115-2021; 2022-2024) and the Czech Ministry of Youth and Education MSCA Fellowships Interfellows (Project number: CZ.02.01.01/00/22_010/0003414; 2024-2025). This work was carried out with the support of the Growth Facility BC Core Facilities, Institute for Plant Molecular Biology, Biology Centre CAS.

## Author contribution

FC and MW conceived and designed the project. FC, AP, and JKW carried out experiments. FC analyzed the data. FC, JKW and MW wrote the article. All authors read and contributed to the final article.

## Data availability

All data is available in the supplementary files provided alongside this manuscript, including: flowering and transcript abundance data, used methodologies, statistical procedures, reagents (including oligonucleotides) and instruments. Sequence data used in this manuscript can be found in the TAIR platform (https://www.arabidopsis.org/) under the following accession nos.: AT1G22690, AT1G15550, AT1G79460, AT5G10140, AT5G08290, AT5G25760.

## Supplementary figure legends

**Figure S1**. The flowering delay of the *crk2* mutant is more severe under SD.

**Figure S2**. Expression of the GA biosynthesis genes *GA2* and *GA3ox1* is altered in *crk2*.

**Figure S3**. The recombinant cytosolic domain of CRK2 (CRK2cyt) phosphorylates recombinant GRP7, but not the GRP7-interactor JACALIN-LECTIN LIKE1 (JAC1), *in vitro*.

**Figure S4**. The *crk2* mutant is sensitive to vernalization but does not show increased *FLC* expression.

## Supplementary table legends

**Table S1**. Photoperiod experiment. Raw data and statistical analysis.

**Table S2**. Flowering time and rosette leaf number at flowering. Raw data and statistical analysis.

**Table S3**. Relative transcript abundance of *GA3ox1 & GASA9*.

**Table S4**. Relative transcript abundance of GA biosynthesis-related genes. Raw data and statistical analysis.

**Table S5**. Vernalization experiment. Normalized data and statistical analysis.

**Table S6**. Relative transcript abundance of FLC. Raw data and statistical analysis.

**Table S7**. List of oligonucleotides used for gene expression analysis.

