## Supplementary information for "CRK2 modulates flowering in *Arabidopsis* together with GLYCINE- RICH RNA BINDING PROTEIN 7 (GRP7)"

**Fig. S1** The flowering delay of the *crk2* mutant is more severe under SD.

**Fig. S2** Expression of the GA biosynthesis genes *GA2* and *GA3ox1* is altered in *crk2*.

**Fig. S3** The recombinant cytosolic domain of CRK2 (CRK2cyt) phosphorylates recombinant GRP7, but not the GRP7-interactor JACALIN-LECTIN LIKE1 (JAC1), *in vitro*..

**Fig. S4** The *crk2* mutant is sensitive to vernalization but does not show increased *FLC* expression.

**Table S1** Photoperiod experiment. Raw data and statistical analysis.

**Table S2** Flowering time and rosette leaf number at flowering. Raw data and statistical analysis.

**Table S3** Relative transcript abundance of *GA3ox1* & *GASA9.*

**Table S4** Relative transcript abundance of GA biosynthesis-related genes. Raw data and statistical analysis.

**Table S5** Vernalization experiment. Normalized data and statistical analysis.

**Table S6** Relative transcript abundance of *FLC*. Raw data and statistical analysis.

**Table S7** List of oligonucleotides used for gene expression analysis.

**Methods S1** Detailed description of the methodologies used in this letter.

**Fig. S1 The flowering delay of the *crk2* mutant is more severe under SD.** Flowering time (in days) of Arabidopsis thaliana Col-0 and *crk2* mutant plants grown under short-day (SD) and long-day (LD) conditions. Data is presented as boxplots showing the median, interquartile range (IQR), and whiskers extending to 1.5×IQR. Individual data points are overlaid. Statistical differences between genotypes within each photoperiod condition were assessed using the Wilcoxon rank-sum test, with p-values adjusted for multiple comparisons using the Benjamini–Hochberg (BH) method. Asterisks indicate significant differences between *crk2* and Col-0 (**p < 0.05).

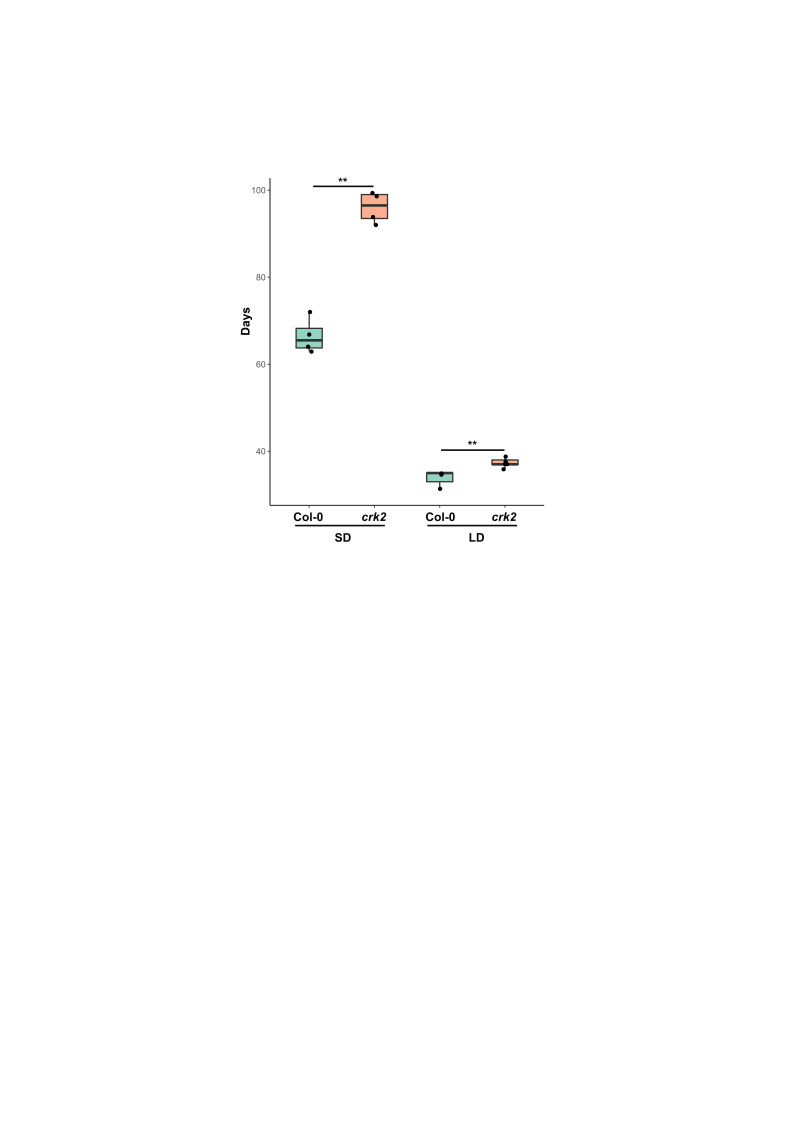

**Fig. S2 Expression of the GA biosynthesis genes *GA2* and *GA3ox1* is altered in *crk2*.** **(a–b)** Relative transcript abundance of *GA2* **(a)** and *GA3ox1* **(b)** in 3- and 4-week-old seedlings respectively. **(a)** Transcript abundance of *GA2* in Col-0 and *crk2* was compared using a two-tailed *t*-test. Asterisks indicate statistically significant differences (** p<0.05). **(b)** Differences among genotypes, namely Col-0, *crk2*, *grp7-1*, the *crk2/grp7-1* double mutant and complementation lines (*crk2/CRK2p::CRK2-YFP* #1-14 and #1-22), were assessed using a non-parametric Kruskal–Wallis test followed by pairwise Wilcoxon rank-sum tests. **(a, b)** Data is presented as boxplots showing the median, interquartile range (IQR), and whiskers extending to 1.5×IQR. Individual data points are overlaid.

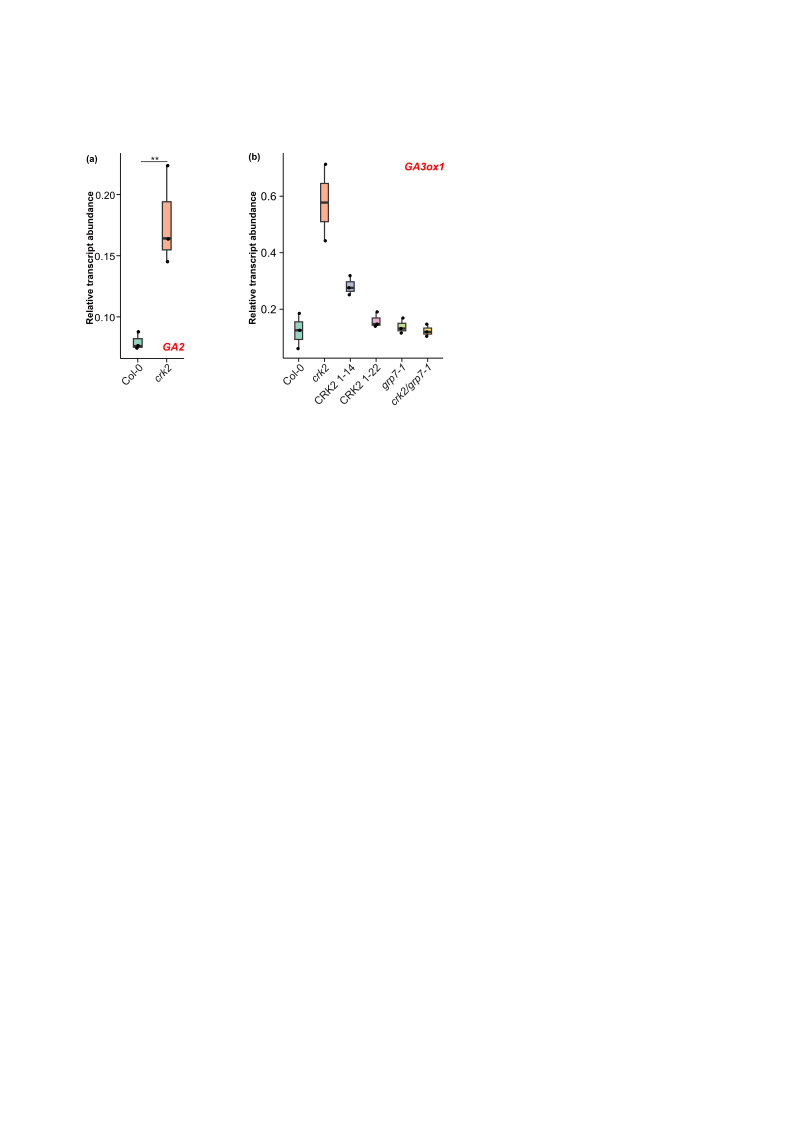

**Fig. S3 The recombinant cytosolic domain of CRK2 (CRK2cyt) phosphorylates recombinant GRP7, but not the GRP7-interactor** **JACALIN-LECTIN LIKE1 (JAC1), *in vitro*.** GST-CRK2cyt Autophosphorylation and transphosphorylation of MBP-GRP7 were visualized with [γ-^32^P] ATP and autoradiography (top). Input proteins were stained with Coomassie Brilliant Blue (CBB; bottom).

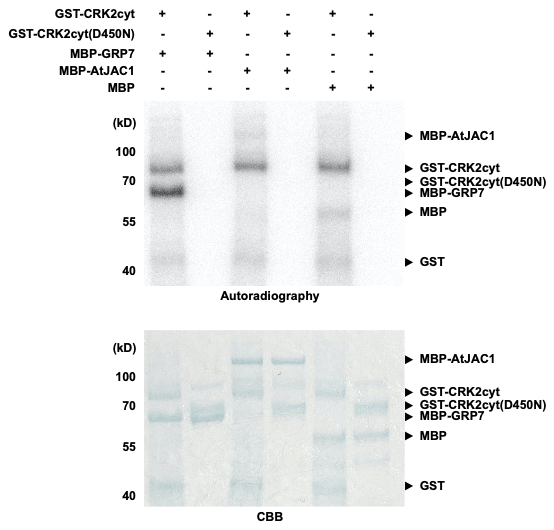

**Fig. S4 The *crk2* mutant is sensitive to vernalization but does not show increased *FLC* expression.** (**a**) Ratio of flowering time (in days) for Col-0, the single mutants *crk2* and *grp7-1*, the *crk2/grp7-1* double mutant and two complementation lines (*crk2/CRK2p::CRK2-YFP*, lines #1-14 and #1-22) with and without vernalization treatment under SD conditions. Pairwise comparisons were performed within each genotype using Wilcoxon rank-sum test. P-values were adjusted for multiple testing using the Benjamini–Hochberg (BH) method (** p<0.05, *** p<0.01). **(b)** Relative transcript abundance of *FLC* in 4-week-old *Arabidopsis thaliana* plants grown under short-day (SD) conditions. Genotypes analyzed included Col-0, the single mutants *crk2* and *grp7-1*, the *crk2/grp7-1* double mutant and two complementation lines (*crk2/CRK2p::CRK2-YFP*, lines #1-14 and #1-22). Differences between genotypes were assessed using a non-parametric Kruskal–Wallis test, followed by pairwise Wilcoxon rank-sum tests. P-values were adjusted for multiple comparisons using the Benjamini–Hochberg (BH) method. Letters indicate statistically significant differences between genotypes (p < 0.05). (**c**) *FLC* expression in Col-0 and *crk2* plants exposed to different cold treatments: no cold (0 d), 12 days, and 45 days. Differences between genotypes were assessed using pairwise Wilcoxon tests with Benjamini–Hochberg correction for multiple comparisons. **(a-c)** Data is presented as boxplots showing the median, interquartile range (IQR), and whiskers extending to 1.5×IQR. Individual data points are overlaid.

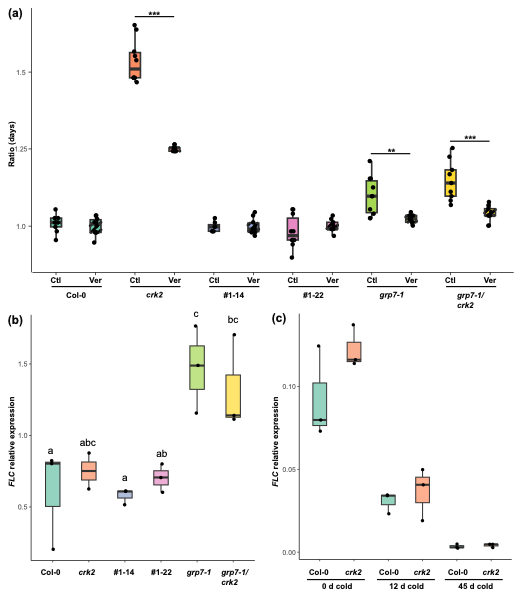

**Table S1 Photoperiod experiment. Raw data and statistical analysis.** Flowering time (in days) of *Arabidopsis thaliana* Col-0 and *crk2* plants grown under contrasting photoperiod conditions: short day (SD) and long day (LD). Normality was tested with the Shapiro–Wilk test, and homogeneity of variances with Levene’s test. Since some groups did not follow these assumptions, pairwise comparisons were performed using the Wilcoxon rank-sum test, with p-values adjusted for multiple comparisons using the Benjamini–Hochberg (BH) method.

**Table S2 Flowering time** **and rosette leaf number at flowering. Raw data and statistical analysis.** Raw flowering data for *Arabidopsis thaliana* genotypes Col-0, *grp7-1*, *crk2*, the *crk2/grp7-1* double mutant, and two complementation lines (*crk2*/CRK2p::CRK2-YFP, lines #1-14 and #1-22). Outliers were identified using the interquartile range (IQR) method. Normality was assessed using the Shapiro–Wilk test and homogeneity of variances using Levene’s test. (a) Flowering time (in days). As parametric assumptions were not fully met, a Kruskal–Wallis test was performed, followed by pairwise Wilcoxon rank-sum tests with Benjamini–Hochberg correction for multiple comparisons. (b) Number of rosette leaves at flowering. As parametric assumptions were met, a one-way ANOVA was performed, followed by Tukey’s honest significant difference (HSD) test for post hoc pairwise comparisons. Significance group letters from post hoc analyses are included in both panels.

**Table S3 Relative transcript abundance of *GA3ox1* & *GASA9*.** Transcript abundance was measured in 2-month-old *Arabidopsis thaliana* plants of the following genotypes: Col-0, *grp7-1*, *crk2*, the *crk2/grp7-1* double mutant, and complementation lines (*crk2*/CRK2p::CRK2-YFP, lines #1-14 and #1-22) under weekly GA or mock treatments. (a) Transcript abundance of *GA3ox1* (2^−ΔCt). (b) Transcript abundance of *GASA9* (2^−ΔCt). Normality was assessed using the Shapiro–Wilk test, and homogeneity of variances with Levene’s test. Since some group pairs did not meet these assumptions, pairwise comparisons were performed using the Wilcoxon rank-sum test. P-values were adjusted for multiple testing using the Benjamini–Hochberg (BH) method.

**Table S4 Relative transcript abundance of GA biosynthesis-related genes. Raw data and statistical** **analysis.** (a) Relative transcript abundance of *GA2* (2^-ΔCt) in 3-week-old Col-0 and *crk2* seedllings. Normality and homogeneity of variances were confirmed using the Shapiro–Wilk and Levene’s tests, respectively. Differences between genotypes were assessed using a two-tailed t-test. (b) Relative transcript abundance (2^-ΔCt) of *GA3ox1* measured in 4-week seedlings from the following genotypes: Col-0, *crk2*, *grp7-1*, the *crk2/grp7-1* double mutant, and complementation lines (*crk2*/CRK2p::CRK2–YFP; #1–14 and #1–22). Since data did not meet parametric assumptions, statistical differences were assessed using pairwise Wilcoxon rank-sum tests. P-values were adjusted for multiple testing using the Benjamini–Hochberg (BH) method (p < 0.1).

**Table S5 Vernalization experiment. Normalized data and statistical analysis.** Normalized data (days to flower) from the vernalization experiment in *Arabidopsis thaliana* genotypes: Col-0, *grp7-1*, *crk2*, the *crk2/grp7-1* double mutant, and complementation lines (*crk2*/CRK2p::CRK2-YFP, lines #1-14 and #1-22). Ratios were calculated by dividing individual values by the average of Col-0 under the corresponding condition (22 °C or 4 °C). Outliers were identified using the interquartile range (IQR) method. Normality was tested with the Shapiro–Wilk test, and homogeneity of variances with Levene’s test. Since some groups did not follow these assumptions, pairwise comparisons were performed using the Wilcoxon rank-sum test, with p-values adjusted for multiple comparisons using the Benjamini–Hochberg (BH) method.

**Table S6 Relative transcript abundance of *FLC*. Raw data and statistical analysis.** (a) Relative transcript abundance of *FLC* (2^-ΔCt ) in 4-week-old *Arabidopsis thaliana* plants of the following genotipes: Col-0, *grp7-1*, *crk2*, the *crk2/grp7-1* double mutant, and complementation lines (*crk2*/CRK2p::CRK2-YFP, lines #1-14 and #1-22). Statistical analysis was performed using one-way ANOVA followed by Tukey’s post hoc test for multiple comparisons. (b) Relative transcript abundance of *FLC* (2^-ΔCt) in 3-week-old Arabidopsis Col-0 and *crk2* plants exposed to different durations of cold treatment (0, 12, or 45 days). Normality was assessed using the Shapiro–Wilk test, and homogeneity of variances with Levene’s test. Since some group pairs did not meet these assumptions, pairwise comparisons were performed using the Wilcoxon rank-sum test. P-values were adjusted for multiple testing using the Benjamini–Hochberg (BH) method.

**Table S7** List of oligonucleotides used for gene expression analysis. AGI identifiers, gene symbols, oligonucleotide sequences (5’–3’), and references are indicated.

**Methods S1** Detailed description of the methodologies used in this letter

**Plant materials and growing conditions**

The Arabidopsis lines used in this study included Col-0, the null single mutants *crk2* (Bourdais *et al.*, 2015), and *grp7-1* (Fu *et al.*, 2007), the *grp7-1/crk2* double mutant and complementation lines (*crk2/CRK2p:CRK2-YFP* #1-14 and #1-22; (Kimura *et al.*, 2020). The double mutant line was generated by crossing the *crk2* and *grp7-1* lines following established protocols with the *crk2* mutant as pollen donor.

For all experiments, seeds were stratified for 2 days at 4 °C. Stratified seeds were germinated in soil (3:1, peat:perlite) for a week under high humidity and SD (8 h light, 23°C /16 h dark, 21°C) before transplanting. Transplanted seedlings were kept under high humidity and SD for an extra week before transferring to specific experiment conditions (50 % humidity).

**Photoperiod experiment**

Photoperiod experiments were conducted under contrasting photoperiod conditions SD (8 h light, 23°C /16 h dark, 21°C) or LD (16 h light, 23°C /8 h dark, 21°C) in 150 μmol m^−2^ s^−1^ photon flux density from warm white LED panels (Photon Systems Instruments LED Fyto-panels). Two plant groups were included: one grown under SD and other grown under LD until flowering. Flowering time was recorded as the number of days from the end of seed stratification until the inflorescence stem reached 1 cm above the rosette.

**Vernalization experiment**

Vernalization experiments were conducted under short-day (SD) conditions, using the same light settings as those described for the photoperiod experiments. Two plant groups were included: one grown continuously at control temperature (23 °C) until flowering, and another subjected to a 45-day vernalization treatment at 4 °C before being transferred to 23 °C to complete development. Flowering time was measured as the number of days from the start of seed stratification until the inflorescence stem reached 1 cm above the rosette. For each genotype, flowering time data was normalized relative to their respective Col-0 controls. To evaluate the effect of vernalization on *FLC* expression, samples were collected from non-vernalized 3-week-old seedlings, as well as from plants at 12 and 45 days after transfer to 4 °C.

**Gibberellin experiment**

Gibberellin treatments were carried out under SD conditions using the same light settings previously described. Two sets of plants were treated once a week with either 100 μM GA₃ (Duchefa Biochemie, G0907.0001) or mock solution. Treatments continued until flowering was imminent in Col-0 plants from the GA-treated group (approximately two months). Both GA and mock solutions were prepared in water containing 1% (v/v) DMSO and 0.02% (v/v) SILWET L-77. On the final day, a last spray treatment was applied, and samples were collected three hours later, at the end of the light period.

**RNA extraction**

Plant material was immediately frozen in liquid nitrogen after harvesting. Frozen material was homogenized with glass beads using a bead mill. Total RNA was extracted using ROTI^®^ZOL (Karl Roth, 9319.2) according to the manufacturer’s instructions but with specific steps dedicated to the Arabidopsis material. Briefly, the RNA was precipitated from the aqueous phase by using a 1:1 mix of isopropanol and a precipitation solution (0.8 M sodium citrate, 1.2 M NaCl). RNA quality was evaluated by agarose gel electrophoresis and quantified using nanodrop spectrophotometer.

**RT-qPCR**

Total RNA was treated with Turbo DNase I included in the TURBO DNA-free™ Kit (Thermo Fisher Scientific, AM1907). First-strand cDNA was synthesized from 0.75 μg of purified total RNA by using the RevertAid cDNA synthesis kit (Thermo Fisher Scientific, K1621). Real-time quantitative PCR (RT–qPCR) reactions were performed using the DB direct PCR SYBR mix SuperSens (DIANA biotechnologies) in an CFX Connect real-time PCR system (Bio-Rad). All kits were used according to the manufacturers’ instructions. Relative quantification of specific mRNA levels was performed using the comparative 2^-ΔCt^ method (Livak & Schmittgen, 2001) by using the gene-specific primers described in Table S7. Expression values were normalized using the Arabidopsis housekeeping gene *YLS8* for *FLC* in the vernalization experiment (Figure S4c, Table S6b), and *PEX4* for *FLC*, *GA2* and *GA3ox1* in all other experiments.

**Statistical analyses**

Statistical analyses were performed using R (version 4.3.2). For all datasets, outliers were identified and removed using the interquartile range (IQR) method. Values below Q1 − 1.5 × IQR or above Q3 + 1.5 × IQR were removed to avoid distortion in subsequent analyses. The assumption of normality was tested using the Shapiro–Wilk test, and homogeneity of variances was assessed using Levene’s test (leveneTest function, car package v. 3.1.3). When these assumptions were met, group differences were evaluated using one-way ANOVA followed by Tukey’s HSD post hoc test. For non-normal and/or heteroskedastic data, the non-parametric Kruskal-Wallis test was used instead, followed by pairwise Wilcoxon rank-sum tests with p-values adjusted using the Benjamini-Hochberg (BH) method (pairwise_wilcox_test function, rstatix package v. 0.7.2). In other cases where only specific comparisons of interest were required, and parametric asumptions were not met, pairwise Wilcoxon tests were applied directly. All statistical visualizations were generated using ggplot2 v. 3.5.2.

**Plasmid construction & protein purification from *E. coli***

MBP-GRP7 and MBP-AtJAC1 constructs for recombinant proteins were generated by classical cloning. The coding regions of GRP7 and AtJAC1 were amplified by PCR and cloned into the pMAL-c2x vector (Addgene, catalog no. 75286). pOPINK-CRK2cyt and pOPINK-CRK2cyt(D450N) constructs, carrying the coding regions of the cytosolic regions of CRK2 (CRK2cyt) and its kinase dead allele CRK2cyt(D450N) were previously described by Kimura *et al.*, 2020.

WT and D450N GST-tagged cytosolic regions of CRK2 were respectively expressed in *Escherichia coli* Shuffle and Lemo21. MBP-tagged GRP7, AtJAC1 (MBP-GRP7, MBP-AtJAC1) and MBP were expressed in *E. coli* BL21. GST-tagged recombinant proteins were purified using glutathione sepharose 4B (GE Healthcare) according to manufacturer’s instructions. MBP-tagged proteins were purified using amylose resin (New England Biolabs) according to manufacturer’s instructions.

***In vitro* kinase assays**

*In vitro* kinase assays were performed following Kimura *et al.*, 2020. Briefly, purified recombinant proteins were incubated with [γ-32P] ATP for 30 min at room temperature in the kinase assay buffer (50 mM HEPES [pH 7.4], 1 mM DTT, 10 mM MgCl2, 0.6 mM unlabeled ATP). The mixture was subsequently separated by SDS-PAGE, and autoradiography was detected by the Typhoon 9410 image analyzer (Amersham Biosciences).
